# Detection and monitoring of translocation renal cell carcinoma via plasma cell-free epigenomic profiling

**DOI:** 10.1101/2025.08.31.673338

**Authors:** Simon Garinet, Karl Semaan, Jiao Li, Ananthan Sadagopan, John Canniff, Noa Phillips, Kelly Klega, Medha Panday, Hunter Savignano, Matthew P. Davidsohn, Kevin Lyons, Alessandro Medda, Prateek Khanna, Mingkee Achom, Prathyusha Konda, Brad J. Fortunato, Rashad Nawfal, Razane El Hajj Chehade, Ze Zhang, Jillian O’Toole, Jack Horst, Dory Freeman, Rachel Trowbridge, Cindy H. Chau, William D. Figg, Jacob E Berchuck, Brian D. Crompton, Ji-Heui Seo, Toni K. Choueiri, Matthew L. Freedman, Sylvan C. Baca, Srinivas R. Viswanathan

## Abstract

*TFE3* translocation renal cell carcinoma (tRCC), an aggressive kidney cancer driven by *TFE3* gene fusions, is frequently misdiagnosed owing to morphologic overlap with other kidney cancer subtypes. Conventional liquid biopsy assays that detect tumor DNA via somatic mutations or copy number alterations are unsuitable for tRCC, since it often lacks recurrent genetic alterations and because fusion breakpoints are highly variable between patients. We reasoned that epigenomic profiling could more effectively detect tRCC, because the driver fusion constitutes an oncogenic transcription factor that alters gene regulation. By defining a TFE3-driven epigenomic signature in tRCC cell lines and detecting it in patient plasma using chromatin immunoprecipitation and sequencing, we distinguished tRCC from clear cell RCC (AUC=0.87) and healthy controls (AUC=0.91) at low tumor fractions (<1%). This work establishes a framework for non-invasive epigenomic detection, diagnosis and monitoring of tRCC, with implications for other mutationally quiet, fusion-driven cancers.

**SIGNIFICANCE:** Translocation renal cell carcinoma (tRCC) is an aggressive fusion-driven subtype of kidney cancer that is frequently misdiagnosed due to morphologic overlap with other kidney cancer subtypes. Conventional liquid biopsy assays targeting DNA alterations are suboptimal for use in tRCC due to its paucity of genomic changes. We demonstrate the utility of cell-free chromatin profiling to noninvasively detect and monitor tRCC with high accuracy, a method that could have applicability to other genomically quiet cancers.

## INTRODUCTION

Translocation renal cell carcinoma (tRCC) is an aggressive subtype of RCC driven by a gene fusion involving an MiT/TFE family transcription factor, most commonly *TFE3*. tRCC comprises up to 5% of renal cell carcinomas in adults (1,2) and over 50% of RCCs in children (3,4). Despite harboring a distinctive genetic landscape, tRCC shares histologic overlap with other RCC subtypes (5), complicating its accurate diagnosis and an estimation of its true incidence. While *TFE3* gene fusions can be detected by break-apart fluorescence in situ hybridization (FISH) (6) or next-generation sequencing (NGS) assays, these tests are not always routinely performed in clinical practice (7–9). Moreover, the variety of *TFE3* fusion partners and genomic breakpoint locations leads to an appreciable false negative rate with these genetic detection methods (10–12). Coupled with the inherent heterogeneity among kidney cancers, with over 40 different subtypes, this creates a critical need for methods that allow for accurate diagnosis and monitoring of tRCC.

Circulating tumor DNA (ctDNA) assays have revolutionized the ability to noninvasively survey molecular features of a patient’s tumor (13–15). To date, most ctDNA assays have focused on the detection of genomic alterations, such as point mutations or copy number variants (16). However, such assays are not suitable for the detection or monitoring of tRCC, which often harbors no genetic alterations apart from the driver fusion (9,17). Moreover, fusion breakpoints are challenging to identify in ctDNA (18), and this is particularly the case in tRCC, where breakpoint locations and *TFE3* partner genes vary widely between patients (9,17,19,20). Indeed, approved cell-free DNA (cfDNA) based liquid biopsy tests do not typically include *TFE3* fusions in the gene panel.

Epigenomic profiling of circulating nucleosomes in plasma has recently emerged as a powerful tool for cancer detection and subtyping that provides orthogonal information to mutation-based methods (21–25). Using an immunoprecipitation-based approach – cell-free ChIP-seq (cf-ChIP) – histone modifications and DNA methylation can be profiled from circulating nucleosomes that originate from cancer cells, enabling the noninvasive measurement of tumor gene regulatory programs in plasma (21,23,26,27).

Given that tRCC is driven by a gene fusion that results in the expression of an oncogenic transcription factor, we hypothesized that cf-ChIP would be uniquely well-suited to detect tRCC-specific profiles of regulatory element activity (28,29), an approach that could also be applied to other fusion-driven malignancies.

## RESULTS

To define an epigenomic signature associated with the TFE3 fusion, we first profiled or reanalyzed 27 epigenomic libraries from 4 tRCC and 6 clear cell RCC (ccRCC) cell lines (**Figure 1 A-B, Methods**). We performed chromatin immunoprecipitation sequencing (ChIP-seq) for two post-translational histone modifications (H3K4me3 and H3K27ac) as well as methylated CpG dinucleotide immunoprecipitation and sequencing (MeDIP-seq). H3K4me3 is enriched at active gene promoters (30) and H3K27ac is enriched at active gene promoters and enhancers (31), while DNA methylation is associated with promoter silencing (32). Across all RCC cell lines, a median of 29,588 peaks (range 25,314-30,076) were captured by H3K4me3 ChIP-seq, a median of 57,226 peaks (44,470-73,400) by H3K27ac ChIP-seq, and a median of 229,624 peaks (111,904-297,344) by MeDIP-seq **(Figure S1)**.

**Figure 1.**
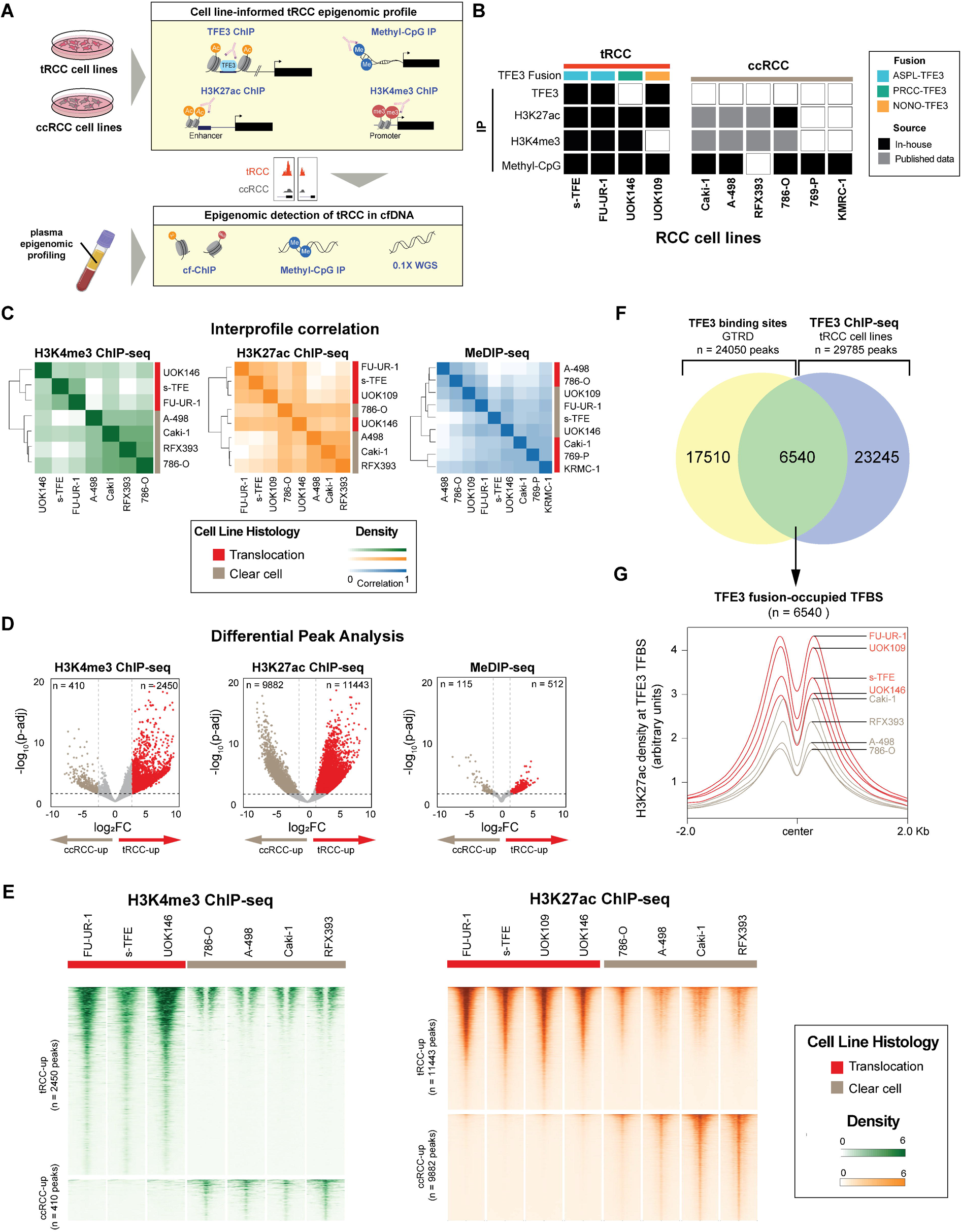
Cell-line informed epigenomic signature of tRCC. **(A)** Overview of study approach: epigenomic profiling of tRCC and ccRCC cell lines was used to inform noninvasive detection of multianalyte epigenomic profiles from plasma cfDNA. **(B)** Epigenomic datasets generated from 4 tRCC (s-TFE, FU-UR-1, UOK109, UOK146) and 6 ccRCC (Caki-1, A-498, RFX393, 786-O, 769-P, KMRC-1) cell lines, either in-house or in a previously published study (61). **(C)** Unsupervised hierarchical clustering of the H3K4me3 ChIP-seq, H3K27ac ChIP-seq and MeDIP-seq consensus peaks across tRCC and ccRCC cell lines analyzed in this study. **(D)** Volcano plots showing differentially marked peaks between tRCC and ccRCC cell lines for H3K4me3 ChIP-Seq, H3K27ac ChIP-seq, and MeDIP-seq. Thresholds for significance were set at FDR-q <0.01 and log_2_fold-change (FC) > 1 for H3K27ac and MeDIP and > 2 for H3K4me3. **(E)** Heatmaps of normalized H3K27ac and H3K4me3 tag densities at differentially marked regions between tRCC and ccRCC cell lines shown over a window ±2 kb from peak center. **(F)** Schema for identifying 6,540 TFE3 fusion-occupied TFBS through the intersection of known TFE3 TFBS (GTRD database) and TFE3 fusion peaks via TFE3 ChIP-seq in tRCC cell lines. **(G)** Aggregated H3K27ac density at 6,540 TFE3 fusion-occupied TFBS across RCC cell lines profiled in this study, showing increased signal in tRCC cell lines (red) compared to ccRCC cell lines (grey).

Unsupervised hierarchical clustering and principal component analyses (PCA) of consensus H3K4me3 and H3K27ac peaks revealed clear segregation of tRCC and ccRCC cell lines, while DNA methylation profiles were not discriminatory between tRCC and ccRCC (**Figure 1C, Figure S1, Methods**). This may be in part because methylation marks reflect mainly cell of origin (which are thought to be identical between ccRCC and tRCC (29,33–35)), while histone marks can provide a dynamic readout of cell state (31), thereby more closely reflecting the transcriptional programs being driven by the TFE3 fusion (9).

We next sought to identify regulatory elements with differential epigenetic modifications in tRCC vs. ccRCC cells. Overlapping peaks across samples were merged for each mark, creating a consensus set of 26,529, 63,322, and 342,285 peaks for the H3K4me3, H3K27ac, and MeDIP profiles, respectively. Via differential peak analysis of ChIP-seq data using the Diffbind R package (36), we identified 2,860 differential H3K4me3 peaks (of which 2,450 were enriched in tRCC; “tRCC-up”; FDR-q<0.01 and Log_2_fold-change (FC) > 2) and 21,325 differential H3K27ac peaks (of which 11,443 were enriched in tRCC; “tRCC-up”; FDR-q<0.01 and Log_2_FC > 1) (**Figure 1D-E; Methods**). In contrast, among MeDIP-seq peaks, we identified only 627 differentially methylated regions (DMRs) between tRCC and ccRCC (FDR-q<0.01 and Log_2_FC > 1, **Figure 1D; Figure S1; Methods**), consistent with the weak segregation observed in unsupervised hierarchical clustering. Motif analysis of the 11,443 tRCC-up H3K27ac peaks identified significant enrichment for sequences bound by TFE3 and its paralog MITF, which share consensus binding sites (37). This suggests that tRCC-up H3K27ac sites include regulatory elements activated by direct binding of TFE3 fusions **(Figure S1)** and is consistent with recent reports that TFE3 fusions may facilitate the organization of enhancer loops (29,38).

Next, we sought to refine our epigenetic signature by incorporating transcriptionally active sites directly bound by the TFE3 fusion in tRCC. First, to obtain a robust consensus set of TFE3 fusion binding sites, we intersected a set of 24,050 wild-type TFE3 transcription factor binding sites (TFBS) derived from two non-RCC cell lines (LoVo, a colorectal cancer cell line, and HepG2, a hepatocellular carcinoma cell line) in the Gene Transcription Regulation Database (GTRD) with 29,785 TFE3 TFBS identified via ChIP-seq in three tRCC cell lines representing two distinct TFE3 fusions (39). This resulted in a final set of 6,540 sites that we deemed “fusion-occupied TFBS” (**Figure 1F; Methods**). Assessing aggregated H3K27ac signal across these 6,540 fusion-occupied TFBS revealed a higher signal in all tRCC cell lines as compared to ccRCC cell lines (**Figure 1G**). Importantly, this difference in signal intensity was less pronounced when considering all TFBS from GTRD (n=24,050) or the fusion non-occupied binding sites which did not overlap with TFE3 fusion ChIP-seq peaks (n=17,510) **(Figure S1),** underscoring the importance of building a robust consensus set of TFE3 fusion-occupied TFBS. Furthermore, only 853 of the 6,540 fusion-occupied TFBS overlapped with the H3K27ac-tRCC-up sites (n=11,443) identified using DiffBind **(Figure S1)**, suggesting that these two methods identify partly non-overlapping regulatory sites associated with tRCC.

Having identified a tRCC-specific epigenomic signature in cell line models, we next evaluated its ability to discriminate plasma from patients with tRCC, ccRCC and healthy controls using cf-ChIP. We profiled 141 epigenomic libraries from 51 plasma samples from patients with tRCC (N=30 samples), ccRCC (N=12), and healthy individuals (N=9) (**Figure 2A; Table S1**). cf-ChIP revealed increased signals at tRCC-up or ccRCC-up peaks in RCC plasma that were absent in plasma from healthy volunteers (**Figure 2B**). For example, in tRCC plasma, H3K4me3 and H3K27ac signals were elevated at the *GPR143* gene locus, a tRCC-specific peak and TFE3-fusion target gene (40). Conversely, we observed an increased H3K4me3 and H3K27ac signal at *C1QL1* gene locus in ccRCC plasma samples. These findings were concordant with the published RNA-seq data in ccRCC and tRCC cell lines, as well as in tRCC and ccRCC tumor samples from a cohort of patients with metastatic ccRCC (41) (**Figure S2**).

**Figure 2.**
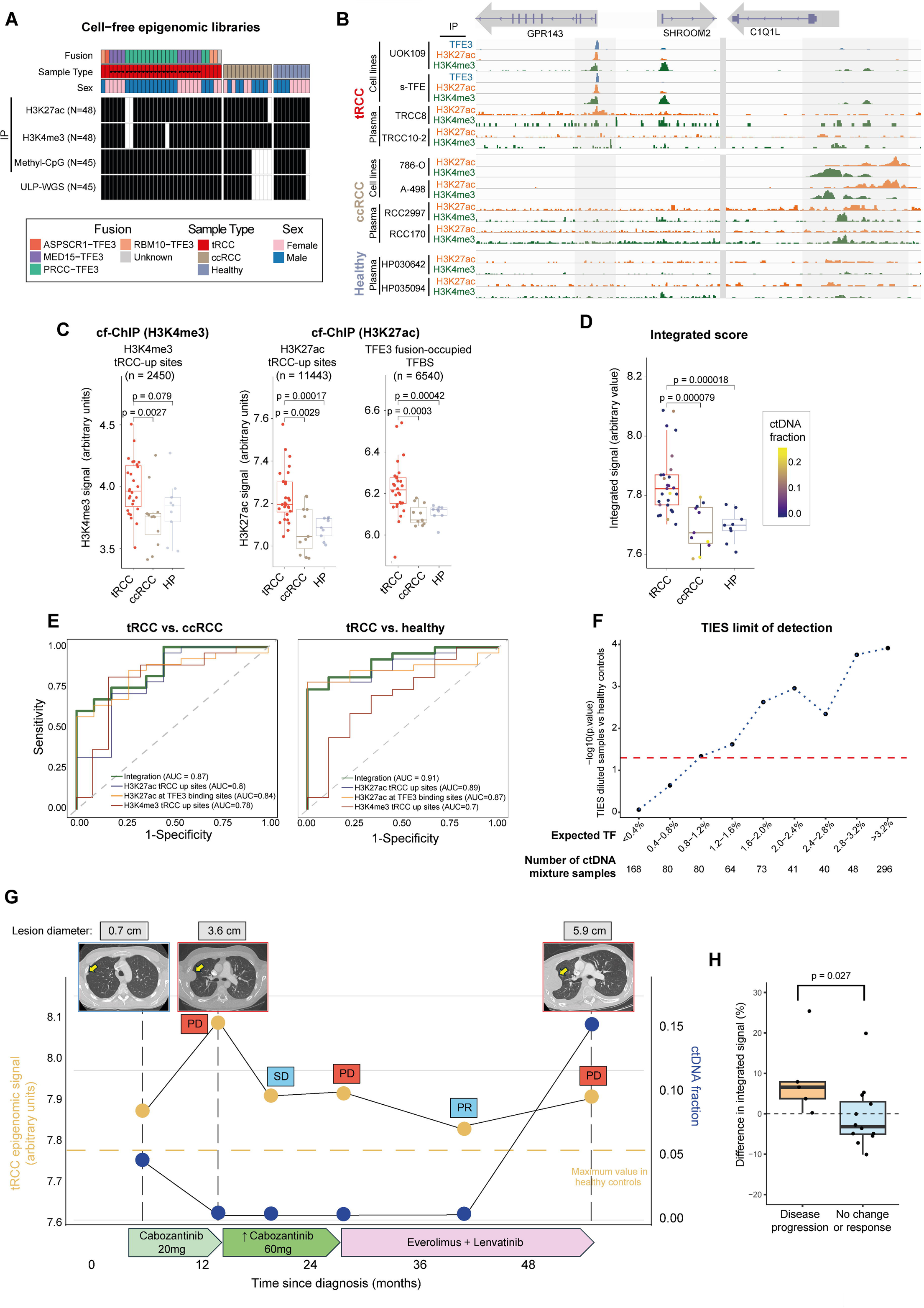
Detection and monitoring of tRCC with plasma cf-ChIP. **(A)** Epigenomic datasets generated from 51 plasma samples collected from patients with metastatic tRCC (N=30; 10 patients), metastatic ccRCC (N=12; 12 patients) or healthy controls (N=9; 9 individuals). **(B)** IGV tracks from ChIP-seq profiles for H3K4me3, H3K27ac and TFE3 in cell lines (tRCC: UOK109, s-TFE; ccRCC: 786-O, A-498) and plasma samples at representative tRCC-selective (*GPR143*) or ccRCC-selective (*C1Q1L*) loci. **(C)** Aggregated cf-ChIP signal compared between tRCC, ccRCC, and healthy plasma samples for the following marks (left to right): H3K4me3 signal at cell line informed H3K4me3 tRCC-up sites (28 tRCC from 10 patients, 11 ccRCC from 11 patients, 9 healthy controls); H3K27ac signal at cell-line-informed H3K27ac tRCC-up sites (27 tRCC from 10 patients, 12 ccRCC from 12 patients, 9 healthy controls from 9 individuals); H3K27ac signal at TFE3 fusion-occupied TFBS (27 tRCC from 10 patients, 12 ccRCC from 12 patients, 9 healthy controls from 9 individuals). The boxplots quantify the area under the curves for each histone mark. **(D)** Comparison of tRCC integrated epigenomic score (TIES) of cf-ChIP H3K4me3 and H3K27ac signals at cell-line informed sites (H3K4me3 tRCC-up peaks, H3K27ac tRCC-up peaks and TFE3 fusion-occupied binding sites), between tRCC, ccRCC and healthy plasma, with samples color-scaled according to tumor fraction (27 tRCC from 10 patients, 11 ccRCC from 11 patients, 9 healthy controls from 9 individuals). **(E)** Classifier assessing individual cf-ChIP H3K4me3 and H3K27ac signals at cell line-informed sites – H3K4me3 tRCC-up peaks, H3K27ac tRCC-up peaks, and TFE3 fusion-occupied binding sites – and evaluating their combined performance in distinguishing tRCC from ccRCC across all patients (left), patients with detectable tumor fraction (>1%) (middle), and tRCC versus healthy plasma (right). **(F)** Estimation of TIES limit of detection. 11 tRCC samples (all with tumor fraction >3% by ichorCNA were diluted *in silico* by adding reads from 8 healthy plasma samples at 0.9, 0.8, 0.7, 0.6, 0.5, 0.4, 0.3, 0.2, 0.1, 0.05, 0.01 ratios. An estimated TF was assigned for each dilution, equal to the TF calculated by ichorCNA for the tRCC sample multiplied by the portion of tRCC reads for that dilution. For a given bin of estimated TF, TIES for all tRCC samples were pooled and compared to healthy plasma value (Wilcoxon sum rank test). Red dashed line indicates the threshold of significance (p=0.05) **(G)** Longitudinal tracking of the tRCC integrated epigenomic score (orange) and ctDNA fraction (blue) in a patient with tRCC. Radiographic changes in an index lesion (pleural metastasis) are provided, as are timings and doses of administered systemic therapies. SD: stable disease; PR: partial response; PD: progressive disease. **(H)** Percentage change in tRCC integrated epigenomic score between consecutive plasma draws, grouped by radiographic response during that same interval.

Given that *GPR143* and *C1QL1* appeared to be highly selective marker genes for tRCC and ccRCC respectively, we also attempted to measure their expression directly in the blood by profiling circulating tumor cells (CTCs). A cohort of 10 patients with metastatic ccRCC, 7 with metastatic tRCC, and one healthy control was sampled for circulating tumor cells (CTCs) using isolation on the TellDx CTC system **(Methods).** We extracted RNA from pelleted samples and aimed to detect tRCC (*GPR143, TRIM63)* or ccRCC (*C1QL1*)-specific transcripts by digital droplet PCR (ddPCR), after whole transcriptome amplification. While these transcripts could be detected in healthy blood spiked with RNA derived from tRCC or ccRCC cell lines, no signal was detectable in the tRCC or ccRCC patient samples. (**Figure S2**). Of note, four patients were sampled for both cfDNA and CTC isolation, with two of them being sampled from the same blood draw. These results suggest that cf-ChIP can infer tRCC gene expression programs even when they are not detectable by standard methods for profiling CTCs.

Next, to develop an epigenome-wide cf-ChIP signature for detecting tRCC, we compared H3K4me3 and H3K27ac coverage in patient plasma at the 2,450 H3K4me3-tRCC-up and 11,443 H3K27ac-tRCC-up peaks, informed by cell line profiling. Aggregated coverage at both H3K4me3 and H3K27ac tRCC-up sites was increased in plasma from patients with tRCC compared to patients with ccRCC (p=0.0027 and p=0.0029, respectively) or healthy controls (p=0.079 and p=0.00017, respectively) **(Figure 2C)**. Conversely, the aggregated methylation signal in plasma at tRCC DMRs, measured using cf-MeDIP-seq, did not distinguish tRCC and ccRCC plasma samples **(Figure S3)**. Finally, H3K27ac coverage at the 6,540 TFE3 fusion-occupied binding sites showed higher discriminating power versus ccRCC (p=0.0003) and versus healthy patients (p=0.00042), perhaps due to wild-type TFE3 activity in white blood cells, particularly macrophages, where MiT/TFE genes can be active (42) **(Figure 2C).**

Finally, we sought to integrate the three sets of epigenomic data described above in order to build a robust cf-ChIP classifier for tRCC. Our tRCC classifier utilized three distinct cell line-informed signatures: H3K4me3 tRCC-up sites, H3K27ac tRCC-up sites, and TFE3 fusion-occupied binding sites. Aggregating plasma H3K4me3 signal across H3K4me3 tRCC-up sites (n=2,450) distinguished tRCC from ccRCC plasma samples (n=27 and n=11, respectively) with an area under the curve (AUC) of 0.78. Aggregating plasma H3K27ac signal across H3K27ac tRCC-up sites (n=11,443) and TFE3 fusion-occupied binding sites (n=6,540) achieved AUCs of 0.8 and 0.84, respectively. To enhance the performance of the classifier, we combined the three distinct scores for each sample, to create a tRCC integrated epigenomic score (TIES). In this process, we ensured that signals at overlapping sites between H3K27ac tRCC-up sites and TFE3 fusion-occupied binding sites were not double-counted **(Figure S1)**. This approach achieved an AUC of 0.87 for the discrimination of tRCC from ccRCC **(Figure 2E)**. For discriminating tRCC from healthy plasma samples (n=27 and n=9, respectively), aggregated H3K4me3 signal at H3K4me3 tRCC-up sites and aggregated H3K27ac signal at H3K27ac tRCC-up sites and TFE3 fusion-occupied binding sites achieved AUCs of 0.7, 0.89 and 0.87, respectively, with an integrated AUC of 0.910 **(Figure 2E)**. Subsequently, we conducted a logistic regression analysis, identifying an actionable TIES cutoff at 7.77. This threshold achieved 100% sensitivity and 71% specificity in detecting tRCC compared to healthy plasma samples. In addition, at single patient level, and among plasma samples with detectable ctDNA, 5 out of 6 patients with tRCC exhibited elevated TIES scores. Interestingly, two tRCC plasma samples had markedly elevated TIES scores but < 3% tumor fraction estimated by ichorCNA, possibly due the paucity of copy number alterations in some tRCC tumors. **(Figure S3**).

Given this observation, we estimated the limit of detection of the TIES via *in silico* dilution. We combined the 11 tRCC samples with tumor fraction > 3% (as estimated by ichorCNA) with each of the 8 healthy plasma samples at 11 dilution ratios (88 combinations for each of 11 dilution levels ranging from 0.9 to 0.01 ratio of tumor:healthy plasma by number of reads). Diluted samples were binned into intervals of 0.4% expected tumor fraction (ranging from <0.4% to >3.2%). In each bin, the TIES score for the tumor dilutions was compared to the TIES score for the 8 healthy plasma samples via Wilcoxon test. A significant difference between healthy samples and tRCC samples (p<0.05) was observed down to a tumor fraction of 0.8-1.2% **(Figure 2F, Figure S3)**, consistent with a limit of detection near 1%.

Having established a tRCC-specific epigenetic signature that can be detected in plasma cfDNA, we also aimed to monitor tRCC disease burden via TIES measured by cf-ChIP in three patients with metastatic tRCC whose plasma samples were collected at multiple time points during treatment **(Figure 2G, Figure S4)**. In all three patients, we observed that variations in TIES were concordant with the clinical course of response and progression to systemic therapy. For instance, in the patient TRCCP4, we observed an increase of TIES at three time points (13-, 26-, and 52-months post-diagnosis) corresponding to radiographic progression but a decrease at time points corresponding to disease control or response (18 months and 39 months post-diagnosis) **(Figure 2G)**. In patient TRCCP5 (initially misdiagnosed as ccRCC on pathology), we observed an increase in TIES at disease recurrence, followed by a subsequent decrease after a change in systemic therapy that resulted in radiographic disease control **(Figure S4)**.

Similarly, in patient TRCCP3, we observed an initial decrease in TIES following curative nephrectomy, followed by an increase of the signal 16 months later, aligned with disease recurrence and metastasis to the liver **(Figure S4)**. We note that, across multiple patients, TIES was detectable and dynamic even when the tumor fraction was in the undetectable range (<3%) by a method that estimates ctDNA using copy number alterations (16). To evaluate cf-ChIP for monitoring tRCC treatment response, we calculated changes in TIES and tumor fraction for each pair of consecutive plasma draws. When we compared these changes during intervals of disease progression, stability, or response, we found that TIES between consecutive draws increased at times of disease progression and decreased during response or disease stability (p=0.027, **Figure 2H**). Importantly, changes in tumor fraction were less pronounced (p=0.63, **Figure S4**). This suggests that our liquid biopsy epigenomic assay may be more effective at tracking disease evolution and response compared to other methods that rely solely on copy number alterations, likely due to the low frequency of such alterations in tRCC.

Finally, to evaluate the extensibility of our approach, we assessed its potential to detect other fusion-specific epigenomic signatures in plasma samples. We compared plasma samples from patients with prostate cancer with (n=5) or without (n=8) the *TMPRSS2-ERG* fusion, previously profiled with cf-ChIP (21). The *TMPRSS2-ERG* fusion, which places the *ETS-*family transcription factor *ERG* under the control of the androgen-responsive gene TMPRSS2, is found in approximately 50% of prostate cancer cases and is associated with a distinctive transcriptional signature (43,44). We observed that plasma from patients with the *TMPRSS2-ERG* fusion exhibited significantly higher H3K27ac signal at fusion-specific H3K27ac sites (n=7,531)(45) compared with samples from patients with fusion-negative cancers (p=0.006); this corresponded to an AUC of 0.95 for discriminating between samples with and without the *TMPRSS2-ERG* fusion (**Figure S5A, S5B**).

We also reasoned that cf-ChIP might be more broadly applicable in cancers with distinctive transcriptional profiles, such as those harboring driver fusions involving a transcription factor. To nominate additional cancer types that may be amenable to profiling via cf-ChIP, we first performed a pan-cancer survey of the fraction of genome altered (FGA), a metric of the proportion of the genome affected by copy number alteration (46). We observed significant variation in FGA both between and within cancer lineages (**Figure S5C**); we note that tumors with low FGA may have insufficient CNAs to enable accurate estimation of tumor fraction in cfDNA (16). We then assessed how FGA tracked with fusion status across cancer types. When limiting to tumors harboring driver fusions involving transcription factors (47,48) (analogous to the *TFE3* fusions in tRCC), we observed that fusion-positive cancers had significantly lower FGA than fusion-negative cancers (median 0.06 vs. 0.20, respectively, p<2.2x10^-16^; **Figure S5D**), consistent with a prior pan-cancer fusion analysis suggesting that >1% cancers may harbor a fusion oncogene as the sole driver (48). For example, FGA in *TFE3* fusion-positive RCCs was significantly lower than in other RCCs (median 0.08 vs. 0.15, respectively, p=0.038). Similarly, FGA in SSX2 fusion-positive synovial sarcoma was significantly lower than in other sarcomas (median 0.08 vs. 0.34, respectively, p=0.011) **(Figure S5E)**. Together, these findings may suggest a potential applicability of cf-ChIP to an array of mutationally quiet cancers, particularly those harboring driver fusions involving a transcription factor.

## DISCUSSION

Plasma epigenomic profiling offers a promising tool for accurate, sensitive, and non-invasive detection of tRCC. Liquid biopsy assays based on tumor genetic changes are challenging to adapt to tRCC, given that it harbors few recurrent genetic mutations and often displays few or no large-scale copy number alterations assessable via low-coverage genome-wide cfDNA sequencing approaches (9,16,17). Another challenge for tRCC detection in plasma is the wide variability of fusion partners and fusion breakpoints (17), which complicates the ability to apply a universal ultrasensitive targeted sequencing approach to this cancer. By contrast, plasma epigenomic profiling enables the inference of tumor-specific regulatory element activity from blood and detection of the TFE3 fusion-driven cistrome in plasma cfDNA regardless of fusion partner or specific breakpoint. Using *in silico* dilution of cf-ChIP reads, we show that tRCC ctDNA can be detected from circulating chromatin at tumor fractions of ∼1%. To our knowledge, this is the first study to employ a non-invasive epigenomic profiling method for accurate detection and monitoring of tRCC in plasma, with potential implications for accurate diagnosis, prognostication and therapy selection. Importantly, this approach can be broadly applied to other cancers that have distinctive epigenomic profiles driven by a transcription factor, but lack sufficient genetic alterations to detect ctDNA with mutation-based methods.

Our findings have several clinical implications. First, underdiagnosis and misdiagnosis of tRCC presents a major challenge, stemming from its histological similarities with other RCC subtypes coupled with the absence of routine molecular testing in clinical practice. Proper diagnosis of tRCC is critical, given that it carries a worse prognosis than other RCC subtypes, may have higher metastatic potential, and may have a lower response rate to systemic therapies typically used for ccRCC (9,49–53). Second, accurate diagnosis of tRCC can help in optimal therapy selection for metastatic disease. While systemic therapies developed for ccRCC are often deployed to patients with tRCC, these therapies typically have lower response rates in tRCC owing to its distinct biology. For instance, belzutifan (HIF-2α inhibitor) has limited mechanistic rationale in tRCC, given that tRCC does not harbor alterations in *VHL* (54). Third, the ability to more accurately detect and diagnose tRCC would have implications for clinical trial enrollment – both to select patients for clinical trials specifically designed for tRCC and to prevent patients with tRCC from being inadvertently enrolled in trials designed for ccRCC, as has occurred with appreciable frequency in the past (9). Finally, we also demonstrate that clinical progression can be preceded by increases in tRCC epigenomic signals detected by cf-ChIP, even at a small systemic disease burden. This assay could therefore improve disease monitoring and inform early treatment-switching. It may also be of use for risk stratification and counseling regarding adjuvant therapy. Adjuvant pembrolizumab following nephrectomy is approved for clear cell RCC but remains unproven and untested in other tumor subtypes like tRCC (55–57), typically being reserved for patients felt to be at highest risk of recurrence and who are most likely to benefit (56).

To date, plasma epigenomic profiling has primarily been used to discriminate tumors of distinct anatomical origin (21,58,59) and cancers with clear histological differences (*e.g.*, neuroendocrine transformation in prostate cancer or lung adenocarcinoma (24,26,60) or sarcomatoid differentiation in RCC (27)). Our study demonstrates that epigenomic profiling of plasma can also accurately distinguish between cancers with overlapping or even indistinguishable histologies, such ccRCC and tRCC. cfChIP-seq is distinctively suited to this purpose because it can detect the activation of regulatory elements by disease-driving transcription factors. Our method overcomes the challenges of detecting gene fusions from ctDNA by measuring the epigenomic effects of TFE3 activation, rather than rearrangement itself. This study has broad applicability to other cancers – particularly those driven by activation of a transcription factor and those that exhibit few copy number alterations and are thus under-detected using DNA alteration-based methods. Our study demonstrates the potential of next-generation liquid biopsy assays that identify transcriptional subtypes of cancer based on epigenomic signatures.

## ACKNOWLEDGEMENTS

S.R.V. acknowledges support from Doris Duke Charitable Foundation (Clinician Scientist Development Award grant number: 2020101); Department of Defense Kidney Cancer Research Program (DoD KCRP) (W81XWH-19-1-0815; W81XWH-22-1-016); Rally Foundation Independent Investigator Award; National Foundation for Cancer Research; V Foundation (V2022-018); NCI (R01CA286652 and R01CA279044); and DF/HCC Kidney SPORE DRP (2P50CA101942-16). SG acknowledges support from ARC (Association pour la Recherche contre le Cancer), la Ligue Contre le Cancer, Institut Servier, Philippe Foundation, and Arthur Sachs grant. P.K. received funding from DoD KCRP Postdoctoral and Clinical Fellowship (HT94252310066

S.C.B. acknowledges support from the US Department of Defense (DoD) award W81XWH-21-1-0299 and the Damon-Runyon Cancer Research Foundation.

M.L.F. acknowledges support from the Claudia Adams Barr Program for Innovative Cancer Research, the H.L. Snyder Medical Research Foundation, the Cutler Family Fund for Prevention and Early Detection, the Donahue Family Fund, the Department of Defense Awards W81XWH-21-1-0234, W81XWH-21-1-0339, W81XWH-19-1-0554, NIH Awards R01CA251555, R01CA227237, R01CA262577, R01CA259058, and a Movember PCF Challenge Award.

T.K.C. is supported in part by the Dana-Farber/Harvard Cancer Center Kidney SPORE (2P50CA101942-16) and Program 5P30CA006516-56, the Kohlberg Chair at Harvard Medical School and the Trust Family, Michael Brigham, Pan Mass Challenge, Hinda and Arthur Marcus Fund and Loker Pinard Funds for Kidney Cancer Research at DFCI.

## AUTHOR CONTRIBUTIONS

S.G., K.S., T.K.C., M.L.F., S.C.B., and S.R.V. conceived the study. S.G. and K.S. led data analysis with assistance from J.L., A.S., B.J.F., P.K, M.A., and M.P. under the joint supervision of S.C.B. and S.R.V. J.C., N.P., H.S., M.P.D., K.L., A.M., P.K., and R.N. performed experiments with input and/or supervision from K.K., J-H.S., B.D.C., M.L.F., S.R.V. J.O’T, J.H., D.F., C.H.C., W.D.F., J.E.B., T.K.C. assisted in procuring clinical samples. S.G., K.S., S.C.B., S.R.V. wrote the first draft of the manuscript and all authors participated in revision of the manuscript.

## METHODS

### Cell lines

786-O (ATCC, CatLog: ATCC® CRL-1932 TM, RRID: CVCL_1051), 293T (ATCC, CatLog: ATCC® CRL-11268TM, RRID: CVCL_0063), A-498 (ATCC, CatLog: ATCC®HTB-44TM, RRID: CVCL_1056), 769-P (ATCC, CatLog: ATCC® CRL-1933 TM, RRID: CVCL_1050), KMRC-1 (JCRB1010, RRID: CVCL_2983), Caki-1(ATCC, CatLog: ATCC® HTB-46TM, RRID: CVCL_0234), UOK109 and UOK146 (Dr. W. Marston Linehan’s laboratory, National Cancer Institute, RRID: CVCL_B123), FU-UR-1(Dr. Masako Ishiguro’s laboratory Fukuoka University School of Medicine) and s-TFE (RIKEN, # RCB4699, RRID: CVCL_6997) were grown at 37°C in DMEM supplemented with 10% FBS, 100 U mL−1 penicillin, and 100 μg mL−1 Normocin (Thermo fisher: #NC9390718).

### Patient cohort

Plasma samples were collected from patients with tRCC (5 males, 5 females) and ccRCC (7 males, 5 females) diagnosed and treated at the Dana-Farber Cancer Institute (DFCI) between 2005 and 2024. All patients provided written informed consent. The collection and use of samples was conducted under an IRB-approved protocol at DFCI. Studies were conducted in accordance with recognized ethical guidelines. Plasma samples from healthy individuals without a history of diabetes, cancer or major medical illnesses were reanalyzed from historical data generated using the same experimental protocol at DFCI, as previously reported (21).

### Sex as a biological variable

In our study, sex was not considered as a biological variable.

### ChIP-seq sample processing

H3K27ac and TFE3 ChIP-seq data in tRCC cell lines that were previously generated in-house (39) are available in GEO under accession number GSE266530. H3K4me3 and H3K27ac ChIP-seq for ccRCC cell lines (Caki-1, A-498, RFX393, and 786-O) were reanalyzed from Gopi et al (61) (GSE143653). H3K4me3 ChIP-seq was performed on tRCC cell lines UOK146, s-TFE and FU-UR-1 as previously described (39). Briefly, 3x10^6^ cells per reaction were collected, crosslinked with 1% formaldehyde and then quenched with 0.125M glycine. After washing with ice-cold PBS, the pellet was resuspended in 130μL SDS Lysis Buffer. Lysate were transferred in sonicated in a Covaris E220 to 200-500bp. Samples were precipitated and centrifuged and 100μL of each sample was diluted 10 times in ChIP dilution buffer and used downstream. 10μL of Dynabeads protein G and 10μL of protein A were washed and complexed with 10μL of H3K4me3 antibody (Rabbit mAb #9751, Cell Signaling, RRID: AB_2616028), and incubated with the lysate at 4°C overnight. Cells were pelleted and washed in 1mL of each of the cold buffers in the order listed below, on a rotating platform at 4°C followed by brief centrifugation and removal of the supernatant fraction on magnetic rack. 1) Low Salt Immune Complex Wash Buffer one wash 5min, 2) High Salt Immune Complex Wash one wash 5min, 3) LiCl Immune Complex Wash Buffer one wash 5min 4) TE, two washes 5min. ChIP DNA was reverse-crosslinked and purified for DNA library construction using the KAPA HyperPrep Kit (KAPA Biosystems, KR0961).

MeDIP-seq was performed on cell lines (UOK146, s-TFE, FU-UR-1, Caki-1, A-498, 786-O, 786-P, and KMRC-1) as previously published described (22). For cfDNA samples, processing for H3K4me3/H3K27ac cf-ChIP and cf-MeDIP-seq on plasma samples was also using previously reported methods (21)(22).

### Analysis of cell line ChIP-seq and MeDIP-seq data

#### Peak calling and identification of tRCC-up regions

For cell line MeDIP-seq and H3K4me3/H3K27ac ChIP-seq data, peak calling was performed using the ChiLin computational pipeline that automates the quality control and data analyses of ChIP-seq (64), with command chilin simple and the parameters –p narrow, -r histone and hg38 as the reference genome. For differential peak analysis between tRCC and ccRCC cell lines, a consensus peak file was generated using Diffbind (v. 3.10.1, RRID: SCR_012918) (36) and an integrated DESeq2-based differential peak calling was performed with DiffBind(36) command dba.analyse. Differentially marked sites were deemed significant if the FDR q-value was < 0.01. We then focused on sites exhibiting a log_2_fold-change in either direction greater than 1 for H3K27ac and greater than 2 for H3K4me3 **(Table S2)**.

#### Identification of fusion-occupied TFE3 transcription factor binding sites (TFBS)

For determination of fusion-occupied TFE3 TFBS, we first downloaded known TFE3 (WT) TFBS from the GTRD database (Homo_sapiens_meta_clusters.zip, http://gtrd.biouml.org:8888/downloads/current/intervals/chip-seq/Homo_sapiens_meta_clusters.zip) (65). This database contains a compilation of ChIP-seq data from various sources with accurate TFBS positions defined by a combination of experiments from various cell lines, and four calling algorithms (MACS2, SISSR, GEM and PICS).

To select TFBS that remained active across TFE3 fusions regardless of fusion partner, we sought to determine a set of high-confidence common peaks between WT TFE3 and various TFE3 fusions. For this, we overlapped the 24,050 peaks available in the GTRD database originating from two non-RCC cell lines (Lovo and HepG2) with 29,785 peaks identified by the union of TFE3 ChIP-seq in tRCC cell lines (FU-UR-1, UOK109 and s-TFE) (39) to determine a set of 6,540 fusion-occupied TFBS. Overlap of peaks was performed using BEDTools v2.27.1 (RRID: SCR_006646)(66). Peaks were considered overlapping if they shared one or more base pairs.

All regions sets were then lifted over from hg38 to hg19 using the UCSC tool “Lift Genome Annotations” (https://genome.ucsc.edu/cgi-bin/hgLiftOver) for comparison with plasma samples **(Table S3)**.

Profile plots of H3K27ac signal at TFE3 TFBS in RCC cell lines were generated using deeptools v 3.5.5 (67) (RRID: SCR_016366)

### Plasma cell-free ChIP-seq

H3K27ac and H3K4me3 cell-free ChIP-seq performed serially on plasma samples using previously published methods (21). The following antibodies were used: H3K27ac, Abcam # ab4729; H3K4me3, Thermo Fisher # PA5-27029.

### Processing and analysis of cfDNA

#### Tumor fraction calculation

cfDNA extraction and low-pass whole-genome sequencing (LPWGS) were performed on plasma samples using previously published methods (21). The ichorCNA R package (RRID:SCR_024768) was used to infer copy-number profiles and cfDNA tumor content from read abundance across bins spanning the genome using default parameters (16). For plasma samples without LPWGS data, we used signal at previously described circulating regulatory elements to estimate cfDNA tumor fraction (21).

#### Analysis of cf-ChIP-seq and cf-MeDIP-seq data

H3K4me3/H3K27ac cf-ChIP-seq and cf-MeDIP-seq reads were aligned to the hg19 human genome build using Burrows–Wheeler Aligner version 0.7.1740 (68) (RRID: SCR_010910). Non-uniquely mapping and redundant reads were discarded. MACS version 2.1.1 (RRID: SCR_013291) (69) was used for ChIP-seq peak calling with a q value (false discovery rate (FDR)) threshold of 0.01. Fragment locations were converted to BED files using BEDTools(66) (version 2.29.2) using bamtobed command with the -bedpe flag set. For analyses involving overlap with genomic regions such a differentially marked regions or TFE3 binding sites, fragments were imported as GRanges objects and collapsed to 1=bp at the center of the fragment location to ensure that a fragment could map to only one site.

#### Quantification of peaks and transcription factor binding sites in plasma

We inferred transcriptional activity at sites of interest based on H3K27ac and H3K4me3, as described previously (21). Briefly, peaks were resized to a 3-kb interval centered on the original peak, then binned into 40-bp windows. Fragment counts were aggregated across each 40-bp window for all peaks to obtain aggregate profiles for each sample. To account for variation in background signal across samples, we performed a ‘shoulder normalization’ step previously described (21). We also normalized signal in each bin to the aggregated signal at the common 10,000 DNAse hypersensitivity sites that are expected to be active across most tissue types and defined across the largest number of samples in reference (70). Similar analysis was used to assess cfDNA methylation patterns at tRCC-DMRs while normalizing signal to the total number of fragments in each sample.

#### Detection of tRCC epigenomic signature in plasma

Our feature set included the differentially marked regions (H3K4me3 and H3K27ac tRCC-up peaks) and TFE3 fusion-occupied binding sites, described above. We measured H3K27ac/H3K4me3 cf-ChIP-seq signal at these sites as described above. The aggregated and normalized signal at these sites was used as an input for the classifier. A subset of 853 peaks, shared between H3K27ac tRCC-up peaks and the TFE3 fusion-occupied binding sites, were discarded from the TFE3 fusion-occupied binding sites when calculating the tRCC epigenomic integrated score to avoid double counting of the signal. The classifier performance was assessed by measuring the area under the receiver operating characteristic (ROC) curve (26), using ROCR R package. In brief, R package the ROC curve was constructed by plotting the true positive rate (sensitivity) against the false positive rate (1 - specificity) for various decision thresholds. The area under the ROC curve (AUC) was then calculated to summarize the overall performance of the classifier.

### Processing of samples for circulating tumor cell isolation

#### CTC Processing

CTCs were isolated from freshly collected whole blood samples as previously described (71). To ensure high recovery of intact CTCs with quality RNA, blood was processed within 4 hours of collection. Leukocytes were depleted using the microfluidic CTC-iChip system (TellBioDX). Whole blood was treated with biotinylated antibodies targeting CD45 clone HI30 (Thermo Fisher Scientific Cat# MHCD4505, RRID:AB_10372216), CD66b clone 80H3 (Standard BioTools Cat# 3162023B, RRID:AB_3661862), and CD16 (BDBiosciences,clone 3G8), followed by incubation with Dynabeads MyOne Streptavidin T1 (Invitrogen) for magnetic labeling and depletion of white blood cells. After enrichment using the CTC-iChip, the CTC product was kept on ice, centrifuged at 4000 rpm, and flash-frozen in liquid nitrogen with RNAlater® (Ambion) for downstream expression analysis.

#### RNA extraction whole transcriptome amplification and ddPCR

RNA was extracted from CTC samples using the RNeasy Plus Micro Kit (Qiagen) following manufacturer’s protocol. Whole transcriptome amplification (WTA) was carried out on RNA samples using the SMARTer® Pico PCR cDNA, following manufacturer’s protocol with 18-21 amplification cycles.

#### Transcript target selection and primer design

Transcripts of interest selective for tRCC or ccRCC were selected based on the analysis of RNA-seq data as described in the manuscript. RNA-seq data for ccRCC cell lines were downloaded from The Cancer Dependency Map Portal (DepMap) (https://depmap.org/portal/download/) RNA-Seq data for tRCC cell lines were previously generated in-house (39) and are under accession number GSE266517. Data for white blood cells were obtained from GTEX V8 (https://gtexportal.org/home/downloads/adult-gtex/bulk_tissue_expression). Transcript targets were selected based on high expression in tRCC or ccRCC (**Figure S2**) and absence of expression in white blood cells defined by <0.5 TPM in GTEX samples. Primers used for ddPCR for specific target genes were obtained from BioRad. *EEF1G*: #10031252-dHsaCPE5191671, *TRIM63*: #10031255-dHsaCPE5049837, *GPR143*: #10031252-dHsaCPE5058198 *C1QL1*: # 10031252-dHsaCPE5052250

#### Digital droplet PCR

cDNA and primer/probe mixes were combined with ddPCR Supermix for Probes (Bio-Rad) in a 96-well plate and loaded into Bio-Rad’s automated droplet generator. Droplets were then subjected to thermal cycling, and droplets containing the target transcripts were detected via fluorescence using the QX200 Droplet Reader System (Bio-Rad).

### cfChIP analysis of *TMPRSS2-ERG* fusion-positive prostate cancer samples

Previously profiled in-house plasma samples from patients with metastatic prostate adenocarcinoma (available in GEO under accession number GSE243474) (21) that had matched panel sequencing data available from the same year as plasma sample were included in the analysis. H3K27ac signal was aggregated at sites differentially marked in fusion-positive compared to fusion-negative prostate cancer identified previously (72), as described in *Quantification of peaks and transcription factor binding sites in plasma* methods section. The classifier performance to distinguish fusion-positive from fusion-negative prostate cancer was assessed by measuring the area under the ROC curve as previously described (26).

### In silico dilution

Cf-ChIP-seq reads from tRCC plasma samples with a tumor fraction > 3% (n=11) and healthy plasma samples with available LP-WGS data (n=8) were randomly selected. We mixed these samples, creating all possible pairwise combinations (n=88) at each dilution level, ranging from 0.9 to 0.01, using Samtools. For each dilution of a tRCC and healthy sample pair, the sample with the fewer reads, S1 (as compared to its counterpart S2), was used to set the read number reference, N, to perform the dilution. At a given dilution level (DL), NxDL reads were randomly sampled from S1 and mixed with Nx(1-DL) reads from S2. The median number of reads combined was 9,331,731 [IQR: 5,228,110-12,071,840].

### TCGA analysis

Fraction genome altered annotations were obtained from the TCGA database for 10,767 tumor samples from 25 cancer types (73) via cBioPortal (74). Driver fusions were taken from a previously published pan-cancer consensus set (75) and limited to those involving transcription factors (47), in analogy to *TFE3* fusions in tRCC. In the pan-cancer analysis, the fraction of genome altered (FGA) was compared between fusion-negative and fusion-positive cancers across all cancer types. In the analysis of *TFE3* fusion in RCC and *SSX2* fusion in synovial sarcoma, we compared fusion-positive RCC and synovial sarcoma with other RCCs and sarcomas without fusions, respectively. Statistical analysis was conducted using the Wilcoxon sum rank test, with a significance level set at 0.05. All analyses were performed using R (v 4.4) on RStudio (v 2024.04.0).

## Data availability

Raw data (FASTQ files) and processed data (BED and BIGWIGS files) are available through GEO under accession numbers GSE280708 (reviewer’s token: ghktiygmjtutpoz), GSE266530, and GSE243474.

## Code availability

Algorithms used for data analysis are all publicly available from the indicated references in the Methods section.

## SUPPLEMENTARY FIGURES

**Figure S1.**
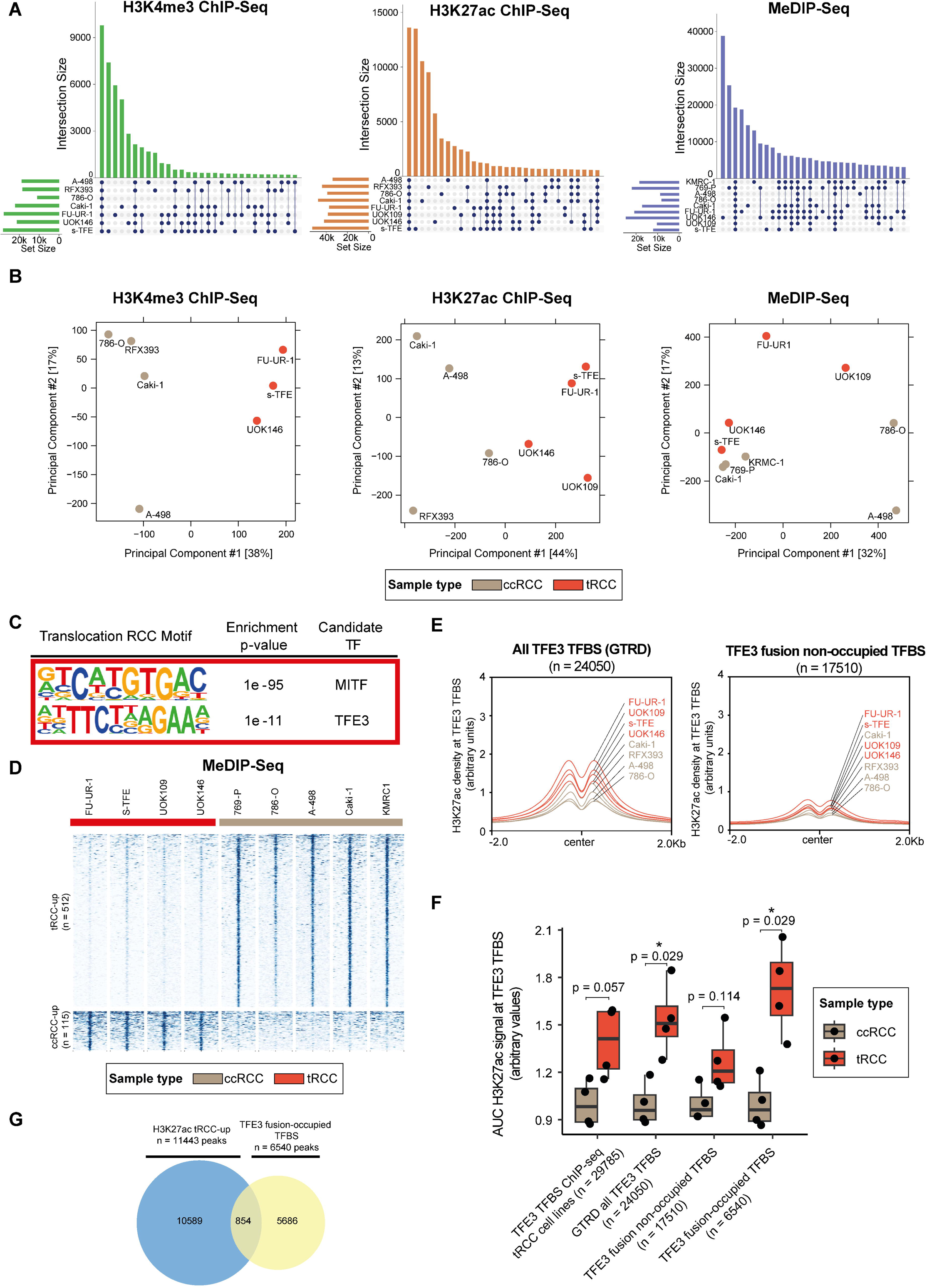
Epigenomic analysis of tRCC and ccRCC cell lines to derive cell-line informed tRCC signature. **(A)** Upset plots showing the intersection of H3K4me3, H3K27ac, and MeDIP peaks across RCC cell lines analyzed in this study. tRCC cell lines: FU-UR-1, UOK146, UOK109, s-TFE; ccRCC cell lines: A-498, RFX393, 786-O, CAKI-1, KMRC-1, 786-P). **(B)** Principal-component analysis (PCA) plots of H3K4me3 ChIP-seq, H3K27ac ChIP-seq and MeDIP-seq peaks for RCC cell lines profiled in this study. **(C)** Motif analysis showing significant enrichment of MITF and TFE3 binding sites amongst 11,443 H3K27ac tRCC-up peaks. **(D)** Heatmaps of normalized MeDIP tag densities at differential MeDIP-seq peak regions between tRCC and ccRCC cell lines (over a window ±2 kb from peak center). **(E)** Aggregated H3K27ac signal density at all 24,050 TFE3 TFBS from GTRD (left) or 17,510 fusion non-occupied TFE3 TFBS in RCC cell lines (right). **(F)** Boxplots of averaged H3K27ac signal at TFE3 TFBS determined respectively from TFE3 ChIP-seq in 3 tRCC cell lines (UOK109, s-TFE, FU-UR-1, N=29,785)), all TFE3 TFBS from GTRD database (N=24,050), TFE3 fusion non-occupied TFBS (N=17,510) or TFE3 fusion-occupied TFBS (N=6,540). P-values were determined by Wilcoxon test. **(G)** Venn diagram showing overlap between H3K27ac tRCC-up peaks (determined by differential analysis of H3K27ac signal in tRCC and ccRCC cell lines) and TFE3 fusion occupied TFBS.

**Figure S2.**
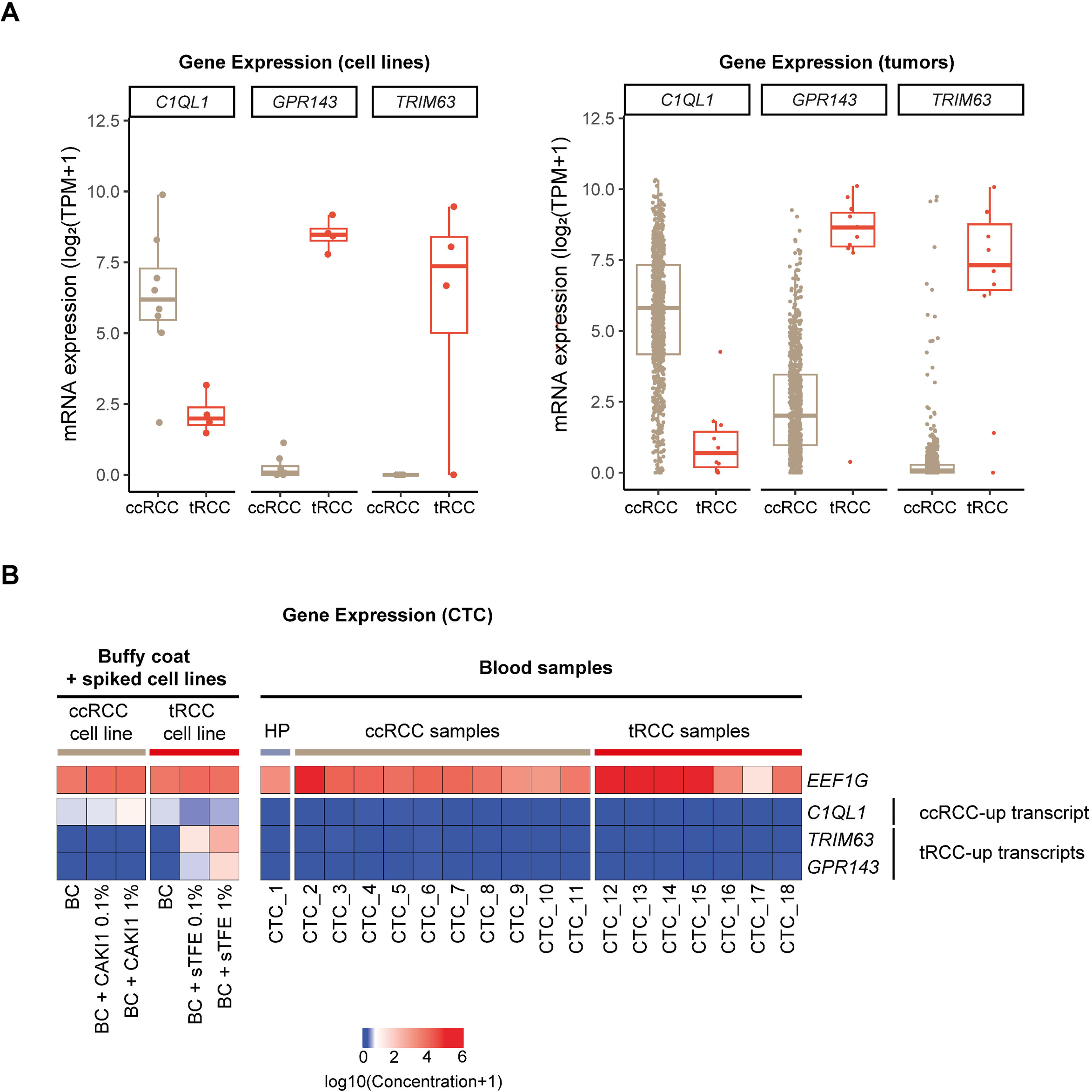
Interrogation of tRCC-specific transcripts in cell lines, tumors, and CTCs. **(A)** Expression of tRCC-selective (*GPR143*, *TRIM63*), and ccRCC-selective (*C1Q1L*) genes in RNA-seq data of RCC cell lines from depmap (tRCC: UOK109, UOK146, s-TFE, FU-UR-1; ccRCC: A-498, ACHN, Caki-1, Caki-2, KMRC-1, KMRC-2, KMRC-20, s786-O) (62) or ccRCC/tRCC tumors from a published study (41). **(B)** Expression of target genes (constitutive: *EEF1G*; ccRCC-selective: *C1QL1*; tRCC-selective: *TRIM63, GPR143*) as detected by ddPCR in CTC isolates from the indicated patients or in healthy blood spiked with either 0.1% or 1% (by RNA quantity) of RNA derived from ccRCC cell line (Caki-1) or tRCC cell line (s-TFE).

**Figure S3.**
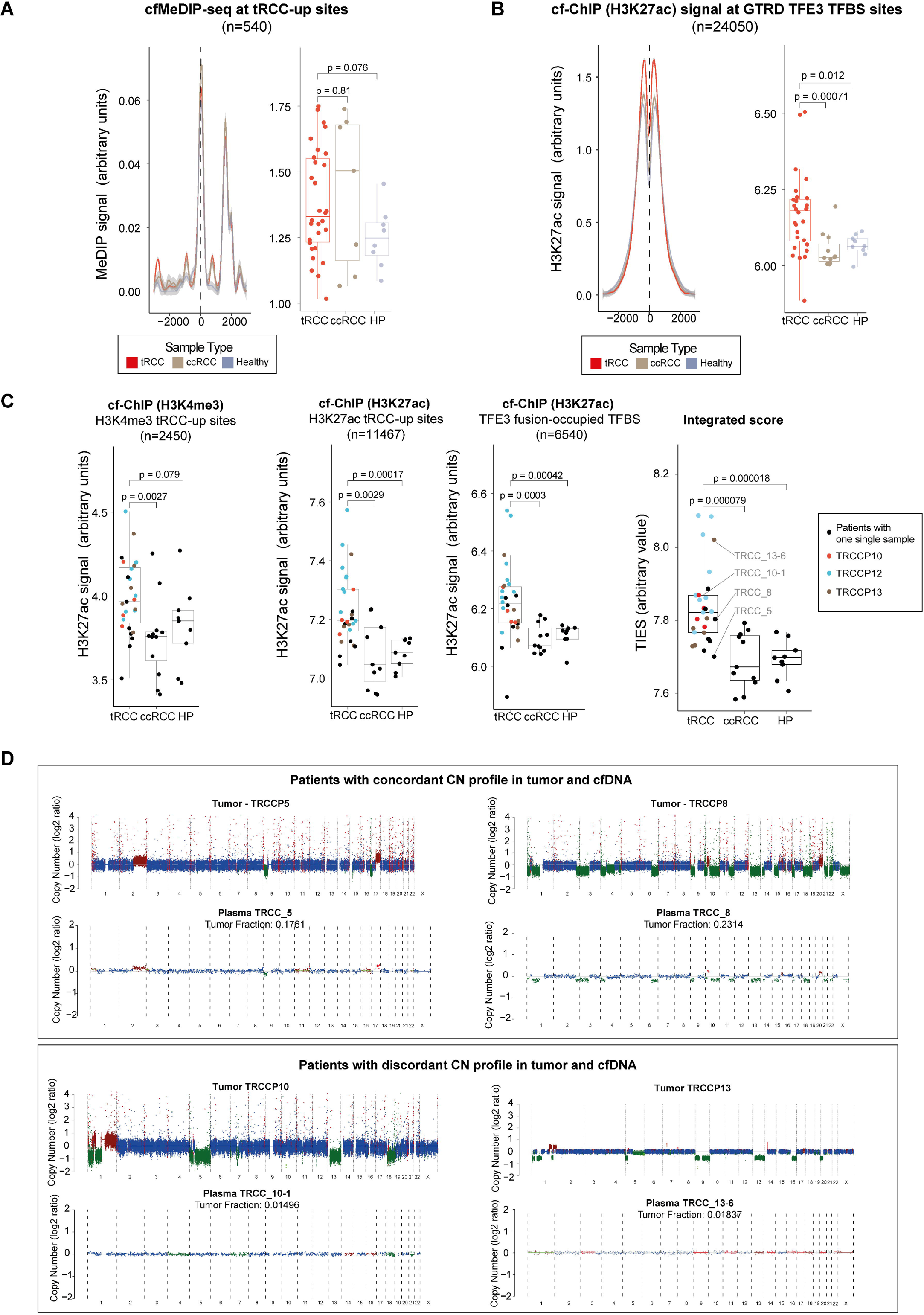
Validation of cf-ChIP in to detect tRCC in plasma. **(A)** Aggregated MeDIP signal at cell-line informed tRCC-up MeDIP sites in tRCC, ccRCC, and healthy plasma samples. **(B)** Aggregated H3K27ac signal at all TFE3 TFBS peaks from GTRD (n=24,050) in tRCC, ccRCC, and healthy plasma samples. **(C)** Aggregated H3K4me3 and H3K27ac cf-ChIP signals at cell-line informed tRCC-up sites (left and middle-left), TFE3 fusion-occupied TFBS (middle-right) and integrated score (TIES, right) across all samples in this study, colored by patient. Samples from patients whose tumors were profiled via WGS (shown in D) are labelled (grey) in the rightmost panel. **(D)** Comparison of genome-wide copy number profiles determined by WGS of plasma cfDNA (ichorCNA (16)) or via WGS of tumors (TITAN (63)), in four patients with matching plasma and samples. Tumor samples were profiled as part of a previously published study (17).

**Figure S4.**
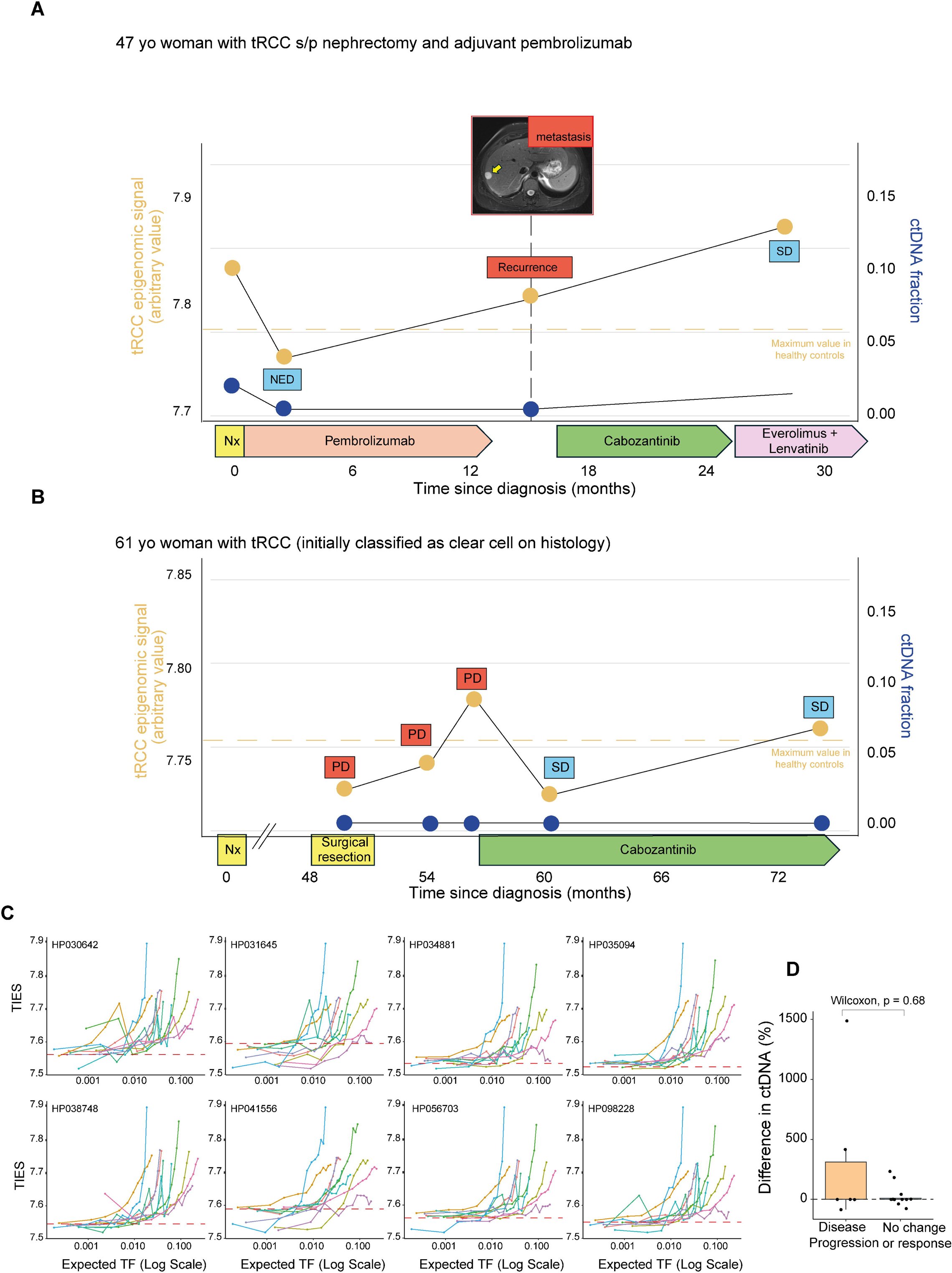
Monitoring of tRCC using cf-ChIP. **(A-B)** Longitudinal tracking of the tRCC integrated epigenomic score (TIES, orange) and ctDNA fraction (blue) in two patients with tRCC. Pertinent radiographic and clinical milestones are indicated in the graphs. SD: stable disease; PR: partial response; PD: progressive disease; NED: no evidence of disease; Nx: nephrectomy. **(C)** *In silico* dilution of tRCC cf-ChIP-seq reads from samples with detectable TF and healthy plasma. TIES was calculated for every pair-wise combination of tRCC and healthy samples at the indicated dilutions. Red dashed line is the TIES of healthy control alone, each tRCC sample is plotted with a distinct color. **(D)** Percentage change in CNA-based cell-free DNA tumor fraction (TF) between consecutive plasma draws grouped by radiographic response during that same interval.

**Figure S5.**
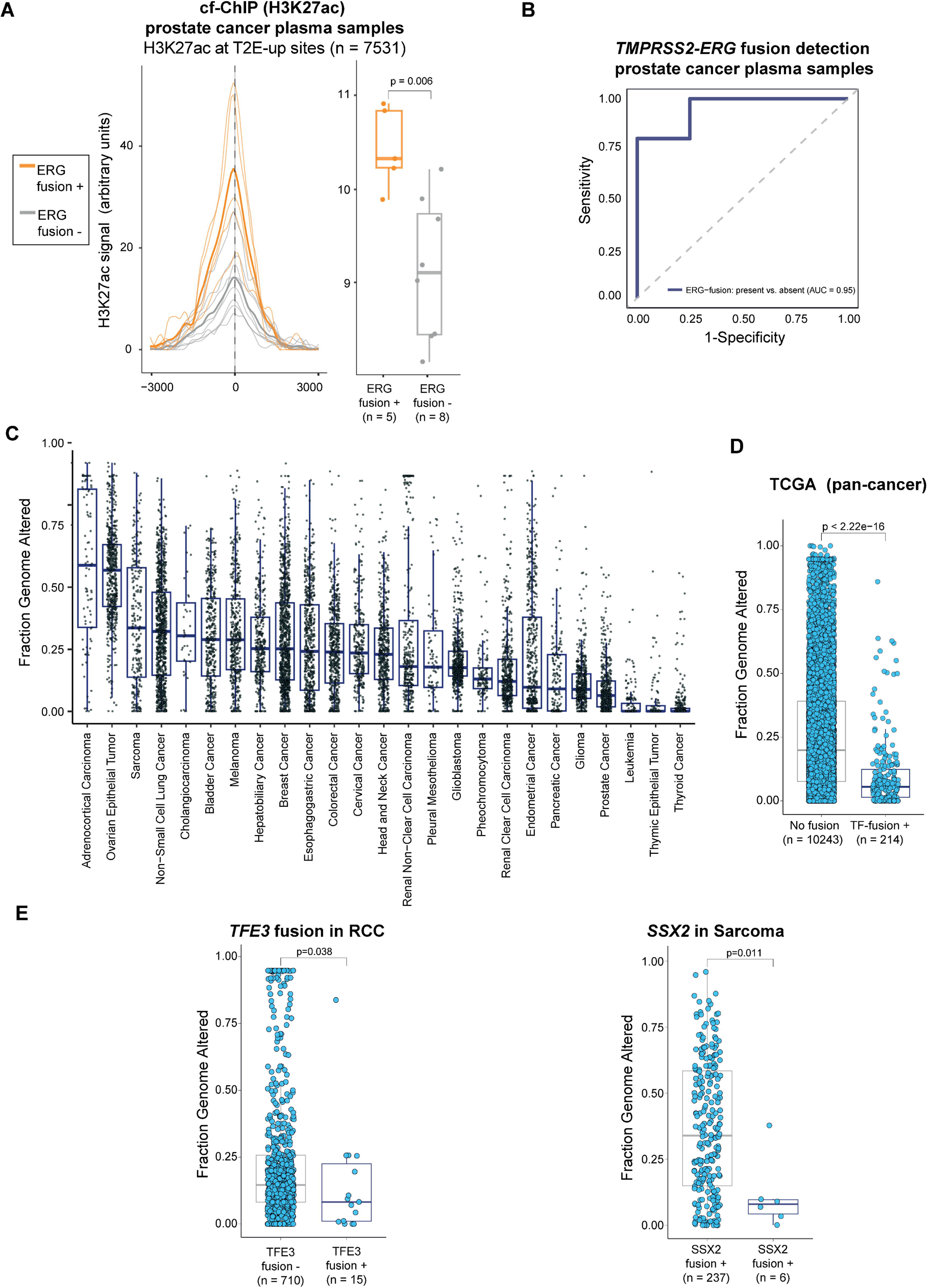
Generalizability of the approach in fusion-driven, mutationally quiet cancers. **(A)** Aggregated cf-ChIP H3K27ac signal compared between *TMPRSS2-ERG* fusion-positive (n=5) and fusion-negative (n=8) prostate cancer plasma samples, at fusion-specific peaks. **(B)** Classifier assessing cf-ChIP H3K27ac signal at *TMPRSS2-ERG* fusion specific peaks in distinguishing fusion-positive and fusion-negative prostate cancer plasma samples. **(C)** Fraction of genome altered (FGA) by cancer type across the TCGA. **(D)** Distribution of FGA in cancers harboring a fusion involving a transcription factor versus fusion-negative cancers, across the TCGA. **(E)** Distribution of FGA in *TFE3* fusion-positive RCC tumor samples vs. other RCCs (left) and SSX2 fusion-positive synovial sarcoma tumor samples vs. other sarcomas (right).

## SUPPLEMENTARY TABLES

**Table S1. Annotation of clinical cohort profiled in this study**

**Table S2. List of H3K27ac and H3K4me3 sites enriched in tRCC (“tRCC-Up”) or ccRCC (“ccRCC-Up”).**

**Table S3. List of consensus fusion occupied TFE3 binding sites used for integrated epigenomic score**

